# TLR9 signaling requires ligand-induced phosphorylation of two specific tyrosine residues by EGFR and Syk

**DOI:** 10.1101/2024.07.03.601759

**Authors:** Manoj Veleeparambil, Chenyao Wang, Patricia M Kessler, Belinda Willard, Ganes C Sen, Saurabh Chattopadhyay

## Abstract

Toll-like receptors (TLRs) are transmembrane proteins required for recognizing microbial components or cellular danger signals to activate intracellular signaling pathways leading to induction of anti-microbial and inflammatory genes. Inactive TLRs require ligand-induced activation to recruit adaptor proteins, e.g., MyD88, to trigger the synthesis of cytokines and interferons. TLR9 is an endosomal membrane-bound protein, which binds to CpG-containing microbial DNA or endogenous signals from dead cells or tissue damage. We showed that TLR9 activation requires EGFR, a tyrosine (Tyr) kinase, which interacts with and phosphorylates the cytoplasmic domain of TLR9. Blocking EGFR activity pharmacologically, or knocking out EGFR gene in myeloid cells, suppressed lethal TLR9-induced hepatotoxicity. Here, we reveal that TLR9 required two Src family of kinases, Syk and Lyn, which, together with EGFR, led to phosphorylation and activation of TLR9. Lack of either of these kinases inhibited TLR9-MyD88 interaction, thereby inhibiting TLR9-mediated gene induction. Unlike EGFR, which constitutively binds TLR9, activated Syk interacted with TLR9 in a CpG-dependent manner. Activated Syk interacted with TLR9 and was critical for activating TLR9-bound EGFR. Quantitative mass spectrometric analyses revealed TLR9 was phosphorylated, sequentially, on Tyr^870^ and Tyr^980^ by Syk and EGFR, respectively. Mutation of either of these tyrosines led to complete loss of TLR9-induced cytokine production. For activation, Syk was phosphorylated by Lyn, which was activated by CpG-mediated scavenger-receptor A, and surprisingly, independent of TLR9. In summary, our results uncovered the molecular details of TLR9 activation by its Tyr-phosphorylation, which is critical for TLR9-mediated intracellular signaling.

## Introduction

Cellular pattern recognition receptors (PRRs), e.g., Toll-like receptors (TLRs), RIG-I-like receptors (RLRs), cyclic GMP-AMP synthase (cGAS) sense viral nucleic acids, also known as pathogen-associated molecular patterns (PAMPs) (1, 2). PRRs also detect cellular danger signals, known as damage-associated molecular patterns (DAMPs), generated from damaged cells and tissues (3, 4). TLRs are transmembrane proteins expressed either on the plasma membrane or endosomal membrane, whereas RLRs and cGAS are localized in the cytosol (5). Specific locations of the PRRs enable them to detect PAMPs from different families of viruses. Upon sensing PAMPs or DAMPs, the PRRs get activated and signal via intracellular adaptor proteins, e.g., TRIF, MyD88, MAVS, and STING, to activate Ser/Thr kinases, e.g., TBK1 and IKKε. Activation of these kinases is required for phosphorylating the inactive transcription factors such as interferon regulatory factor 3 (Irf3) (6). Phosphorylation of Irf3 causes it to dimerize and translocate into the nucleus, leading to the transcriptionally active Irf3, which binds the specific gene promoters to induce antiviral genes, e.g., interferons (IFNs) and IFN-stimulated genes (ISGs). Many of these newly induced ISGs act as viral restriction factors by interfering with specific stages of the viral life cycle to inhibit viral replication (7, 8). PRRs activate, in addition to Irf3, NF-κB, a pro-inflammatory transcription factor, which is responsible for inflammatory gene expression (2, 9-11). Cooperative action of IRF3 and NF-κB is essential for the synthesis of IFNβ, an antiviral cytokine. IFNβ, after synthesized in the virus-infected cells, gets secreted and acts on infected or neighboring uninfected cells to amplify the expression of antiviral genes (12, 13). Therefore, PRR activation is essential for the successful outcomes of the viral infection and inflammatory responses of the host.

TLR9, a member of the TLR family, expressed primarily in immune cells, e.g., dendritic cells, macrophages, as well as B cells, is an endosomal membrane-bound TLR. TLR9 detects the unmethylated CpG containing DNA from PAMPs and DAMPs. Recent studies indicated that TLR9 can also recognize mitochondrial DNA (mtDNA), released from diseased tissues or damaged cells (14, 15). Inactive TLR9 remains as a monomer, which, upon binding CpG DNA by its ectodomain in the endosomal lumen, dimerizes and changes its conformation. The conformational change allows the cytoplasmic domain of TLR9 to recruit its adaptor protein, MyD88. TLR9/MyD88 complex triggers a cascade of intracellular signaling pathways via TRAF6 and IKK to activate the NF-κB. Activation of NF-κB leads to the induction of numerous pro-inflammatory genes, which are essential for regulating cellular inflammatory responses. TLR9 also activates IRF7 via phosphorylation by IRAK1 to induce type-I IFNs (16, 17). UNC93B1, a cofactor for other endosomal TLRs, is involved in TLR9 processing, which is critical for its localization and downstream signaling (18-20). The physiological relevance of TLR9 functions has been established in both viral and bacterial infection models. In addition, the TLR9 ligand, CpG, used as an adjuvant for vaccines, has implications in B cell functions. A plethora of studies have connected TLR9 with non-microbial diseases, e.g., hepatitis and autoimmune diseases (21-23). In such scenarios, TLR9 activation is caused by mtDNA released from dead cells.

TLRs require post-translational modifications, e.g., Tyr phosphorylation of their cytoplasmic domains, for triggering intracellular signaling pathways (24-27). We showed, using extensive biochemical and genetic approaches, the role of TLR3 Tyr phosphorylation in its antiviral and pro-inflammatory functions (27-29). Phosphorylation of TLR3 is critical for recruiting its adaptor protein, TRIF. EGFR and Src phosphorylate two Tyr residues of TLR3 cytoplasmic domain; mutation of either of these two residues results in impaired TLR3 signaling (29). We expanded the role of EGFR in TLR9 phosphorylation and intracellular signaling. EGFR remains bound constitutively to TLR9, and ligation with CpG leads to EGFR-mediated phosphorylation of TLR9 (30). Phosphorylated TLR9 is critical for interaction with its adaptor protein, MyD88, and activating downstream signaling. EGFR inhibitors, often used clinically to treat cancer, block TLR9 signaling *in vitro* and also protect mice from TLR9-induced hepatotoxicity. The role of EGFR-mediated Tyr phosphorylation is known in TLR9 functions (30). However, the molecular details of TLR9 activation leading to its Tyr phosphorylation are incompletely understood. In the current study, we revealed the step-wise activation mechanism of TLR9 by EGFR and Syk-mediated phosphorylation of two Tyr residues in the TLR9 cytoplasmic domain. We took a combination of biochemical, genetic and proteomic approaches to uncover the early events of TLR9 signaling that shapes the eventual outcome of TLR9-mediated cellular responses.

## Results

### Lyn and Syk, in addition to EGFR, are required for TLR9-MyD88 interaction and TLR9-mediated gene induction

We showed that EGFR activity is essential for TLR9-mediated gene induction (30). To evaluate the role of SFKs, particularly Lyn and Syk, in TLR9-induced genes, we generated knockdown (KD) HEK293XL (HEK)-expressing TLR9 (HEK-TLR9) cell lines (EKD: EGFR KD, SKD: Syk KD, and LKD: Lyn KD) by stably expressing kinase-specific shRNAs. Expression of specific shRNA, as expected, did not alter the expression of other Tyr kinases or HA-TLR9 (Fig 1A). Using these cells, we evaluated TLR9-induced gene expression. EKD cells, as expected, did not induce TNF or IFNB1 upon CpG treatment compared to the control cells, expressing non-targeting (NT) shRNA (Fig 1B). Both SKD and LKD cells, like EKD cells, also failed to induce TNF and IFNB1 upon CpG treatment (Fig 1C, D), indicating functional requirements of Lyn and Syk in TLR9-mediated gene induction. TLR9 interacts with its adaptor, MyD88, in a CpG-dependent manner for gene induction, and EGFR is essential for TLR9-MyD88 complex formation (30). We evaluated whether Lyn and Syk were required for TLR9-MyD88 interaction. Indeed, co-immunoprecipitation (co-IP) results indicated TLR9-MyD88 interaction was inhibited in SKD, LKD cells, and, expectedly, in EKD cells (Fig 2A), suggesting these Tyr kinases were essential for TLR9-MyD88 complex formation. We validated these results using R406, a Syk kinase inhibitor, which, as expected, also inhibited TLR9-MyD88 interaction (Fig 2B). Together, EGFR, Lyn, and Syk were required for TLR9-MyD88 interaction and, subsequently, TLR9-mediated gene induction.

**Fig 1.**
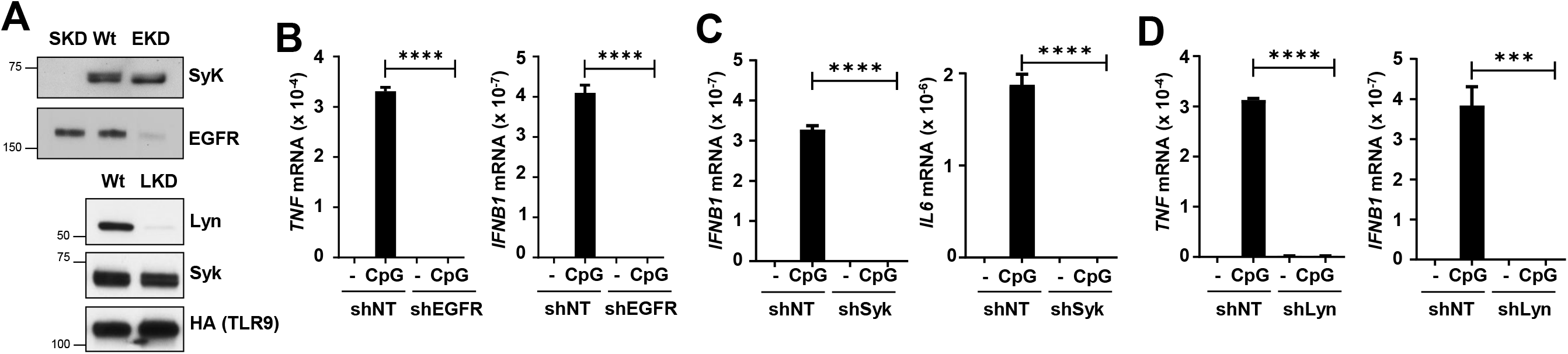
EGFR, Lyn, and Syk are required for TLR9-induced genes. **A)** EGFR, Syk, Lyn expression levels in shNT (Wt) and different gene knockdown 293XL-hTLR9-HA cells were analyzed by immunoblot. **B)** 293XL-hTLR9-HA cells expressing non-targeting shRNA control (shNT) and shRNA targeting EGFR (shEGFR) were treated with CpG ODN (10 μg/ml) for 6 hr and the induced TNF and IFNB1 mRNAs were measured by qRT-PCR. **C)** 293XL-hTLR9-HA expressing shRNA targeting Syk (shSyk) and shNT were treated as in ‘B’ and IFNB1 and IL6 mRNAs were measured by qRT-PCR. **D)** ShNT and 293XL-hTLR9-HA expressing shRNA targeting Lyn (shLyn) were used to measure TNF and IFNB1 mRNA induction by qRT-PCR.

**Fig 2.**
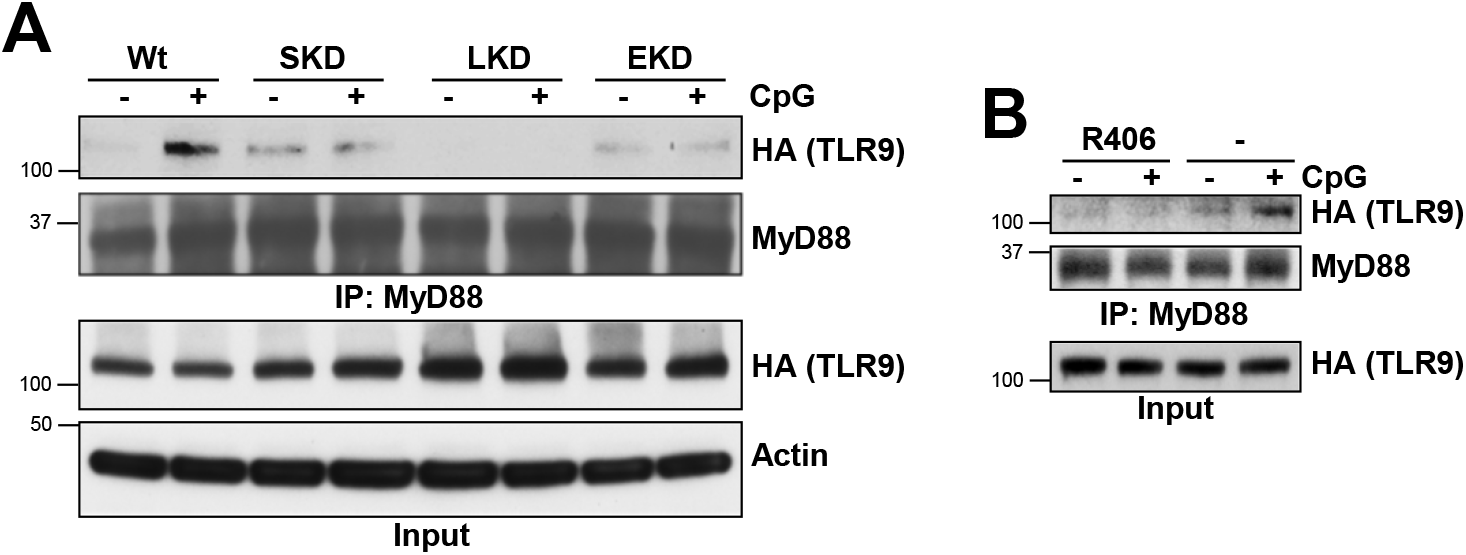
Ligand-induced MyD88 recruitment by TLR9 requires EGFR, Lyn, and Syk. **A)** Wt, shSyk (SKD), shLyn (LKD), and shEGFR (EKD) 293XL-hTLR9-HA cells were treated with CpG ODN for 30 min. Cell extracts were immunoprecipitated with anti-MyD88 followed by immunoblot with anti-MyD88 and anti-HA for TLR9. **B)** 293XL-hTLR9-HA cells were pretreated with DMSO (-) or the Syk kinase inhibitor (R406) (10μM) for 1 hr and then for 30 min CpG DNA along with DMSO or R406. Cell extracts were immunoprecipitated with anti-MyD88 followed by immunoblot with anti-MyD88 and anti-HA for TLR9.

### TLR9 interacts with activated Syk

To investigate whether Syk gets recruited to TLR9 signaling complex, we performed co-IP studies. CpG-stimulation caused rapid recruitment of phosphorylated Syk (pSyk) to TLR9 (Fig 3A). R406 treatment reduced Syk interaction with TLR9 (Fig 3B), indicating activation of Syk was required for TLR9 interaction. TLR9-Syk complex was transient; Syk was recruited to TLR9 in the early phase (5 and 15 min) but dissociated from TLR9 30 min post-CpG-treatment (Fig 3A). TLR9-Syk complex, however, was stabilized in EKD cells, compared to Wt, LKD, or SKD cells 30 min post-CpG-treatment (Fig 3C), suggesting dissociation of Syk from TLR9 was EGFR-dependent. EGFR, however, was not required for recruitment of Syk to TLR9; EKD cells also formed TLR9-Syk complex upon CpG treatment (Fig 3D). Syk and Lyn, on the other hand, did not affect TLR9-EGFR binding; in CpG-treated cells, TLR9-EGFR interaction was unaffected in SKD and LKD cells (Fig 3E). TLR9-bound EGFR was active, analyzed by phosphorylation of its cytoplasmic domain (Y^1068^), only in Wt but not in SKD or LKD cells (Fig 3E), indicating that Syk and Lyn, although not required for TLR9-EGFR interaction, were required for activation of TLR9-bound EGFR. Therefore, EGFR regulated TLR9-Syk interaction; however, although Syk and Lyn had no role in TLR9-EGFR interaction, they were required for activating TLR9-bound EGFR.

**Fig 3.**
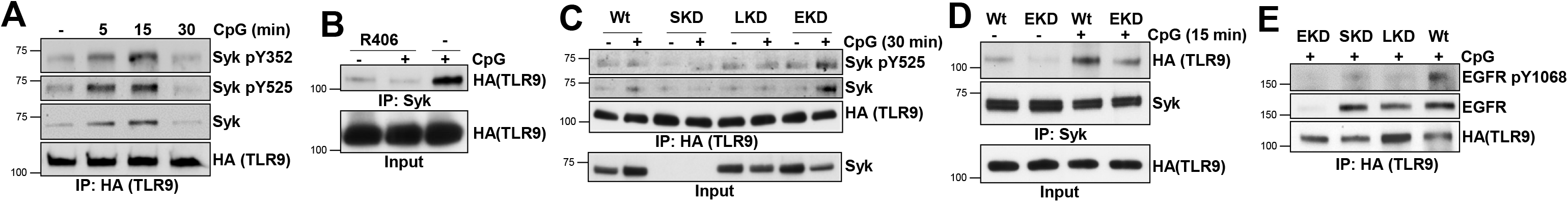
Activated Syk interacts with TLR9 transiently, EGFR inhibits the stability of TLR9-Syk complex. **A)** 293XL-hTLR9-HA cells were treated with CpG ODN for the indicated time, and cell extracts were immunoprecipitated with anti-HA for TLR9 and immunoblotted with the indicated antibodies. **B)** 293XL-hTLR9-HA cells were pretreated with DMSO (-) or the Syk inhibitor (R406) (10 μM) for 1 hr and then treated with CpG ODN along with DMSO or R406 for 15 min. Cell extracts were immunoprecipitated with anti-Syk and immunoblotted with anti-HA for TLR9. **C)** ShNT (Wt), shSyk (SKD), shLyn (LKD) and shEGFR (EKD) 293XL-hTLR9-HA cells were treated with CpG ODN for 30 min, and the cell extracts were immunoprecipitated with anti-HA for TLR9 and immunoblotted with the indicated antibodies. **D)** 293XL-hTLR9-HA shNT cells (Wt) and EKD cells were treated with CpG ODN for 15 min, cell extracts were immunoprecipitated with anti-Syk and immunoblotted with anti-HA for TLR9 and anti-Syk. **E)** Cells and treatment as in ‘D’, cell extracts were immunoprecipitated with anti-HA for TLR9 and immunoblotted with the indicated antibodies.

### The cytoplasmic domain of endosomal EGFR was sufficient for TLR9 signaling

We enquired whether the ectodomain of EGFR, which binds EGF for its activation, was required for TLR9 signaling. To address this, we expressed a membrane-bound form of extracellular domain-deleted EGFR mutant (ΔE) in EGFR KO cells. The ΔE-expressing cells were able to induce TLR9-dependent genes, which were also suppressed by inhibitors of EGFR (Gf) or Syk (R406) (Fig 4A). Since the cytoplasmic domain of EGFR was sufficient for TLR9-mediated gene induction, we evaluated whether ΔE binds TLR9. Indeed, the ΔE mutant, like the full-length EGFR, interacted with TLR9 either in the absence or the presence of CpG (Fig 4B). We validated these results using confocal microscopy, which indicated that ΔE mutant co-localized with TLR9 (Fig 4C). Furthermore, the TLR9-EGFR complex was endosomal, indicated by its colocalization with EEA1, an endosomal marker (Fig 4C). Syk was required for CpG-mediated activation of full-length EGFR (Fig 3E); ΔE mutant, which interacted with TLR9, was, as expected, inactive in the presence of Syk inhibitor, R406 (Fig 4D). Endosomal acidification is a pre-requisite for TLR9-dependent signaling (31); TLR9-mediated gene induction was inhibited by chloroquine (CQ), an inhibitor of endosomal acidification, in both Wt EGFR and ΔE-expressing cells (Fig 4E). Collectively, our results indicated that the cytoplasmic domain of EGFR, which interacted with TLR9, was sufficient for TLR9-mediated gene induction. Furthermore, EGFR ectodomain, which is essential for EGF signaling, was dispensable for TLR9 functions.

**Fig 4.**
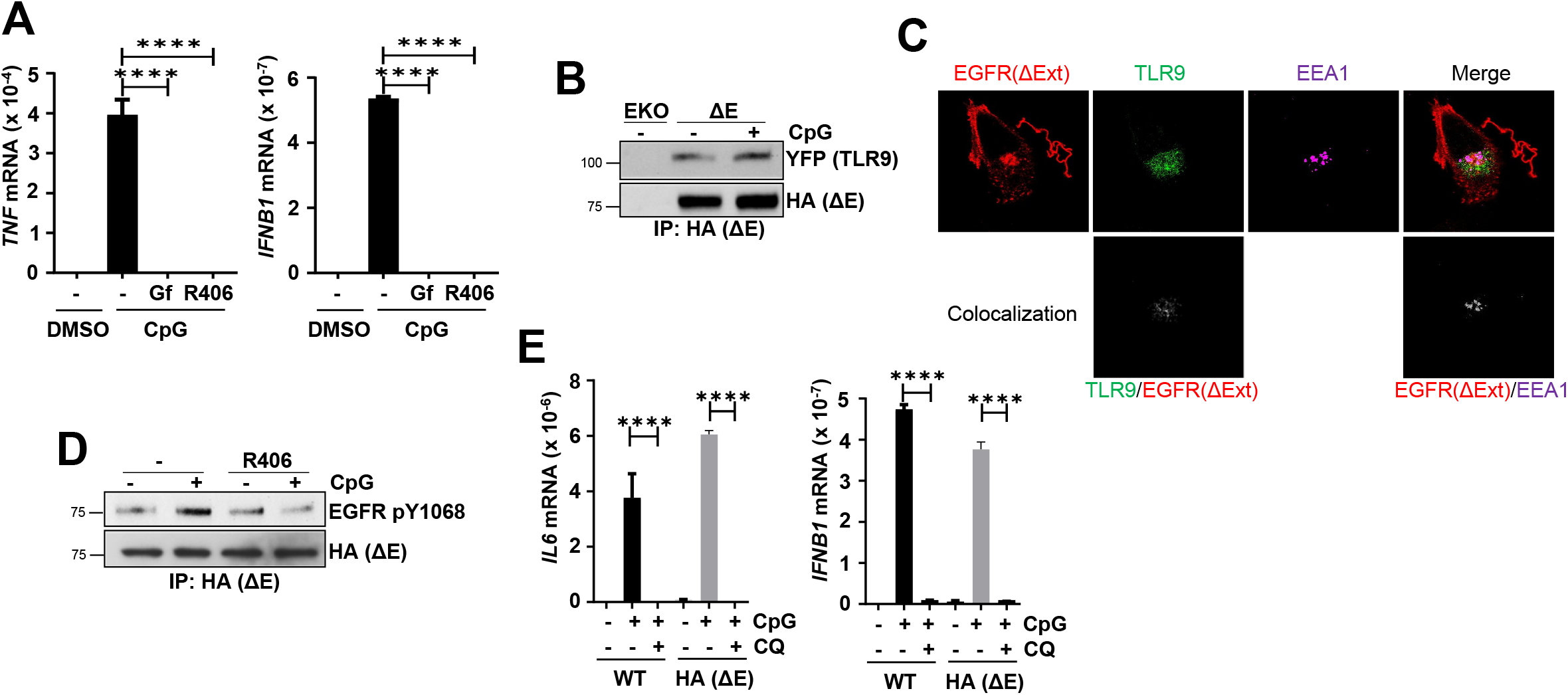
The ligand-binding extracellular domain of EGFR is not needed for TLR9-mediated gene induction. **A)** hTLR9-HT1080 EGFRKO cells reconstituted with extracellular domain deleted EGFR (EGFRΔExt-HA, HA (ΔE)) were pre-treated with DMSO (-) or the EGFR inhibitor, gefitinib (Gf) (10 μM) or Syk inhibitor (R406) (10 μM) for 1 hr and then treated with CpG ODN along with DMSO or Gf or R406 for 6 hrs and TNF and IFNB1 mRNAs were measured by qRT-PCR. **B)** HT1080 EGFRKO cells (EKO) and EKO cells reconstituted with EGFRΔExt-HA and hTRL9-YFP-Flag (ΔE) were treated with CpG ODN for 15 min, cell extracts were immunoprecipitated with anti-HA for EGFRΔExt (HA (ΔE)) and immunoblotted with anti-YFP for TLR9 and anti-HA for EGFRΔExt (HA (ΔE)). **C)** Untreated HT1080 EGFRKO cells reconstituted with EGFRΔExt and hTRL9-YFP-flag were used for immunofluorescence. Red is EGFRΔExt, green is for TLR9, and magenta is for early endosomal marker, EEA1. The bottom two panels are co-localization images using ImageJ co-localization plugin; the white dots represent co-localization. First image is for TLR9/EGFR ΔExt interaction, and the second is for EGFRΔExt/EEA1 interaction. **D)** Cells, as in ‘C’, were pretreated with DMSO (-) or R406 for 1 hr, followed by CpG ODN along with DMSO or R406 for 30 min, cell extracts were immunoprecipitated with anti-HA for EGFRΔExt (HA (ΔE)) and immunoblotted with anti-EGFR pY1068 and anti-HA for EGFRΔExt (HA (ΔE)). **E)** WT and EGFRΔExt (HA (ΔE)) cells were pretreated with chloroquine (CQ) (100 μM) for 1 hr before CpG ODN treatment along with CQ for 6 hr; IL6 and IFNB1 mRNAs were measured by qRT-PCR.

### Mass spectrometric analyses revealed TLR9 is phosphorylated on Y^870^ and Y^980^ by Syk and EGFR, respectively

Given the essential role of TLR9 Tyr phosphorylation in its activation, we determined the specific Tyr residues phosphorylated upon CpG treatment. TLR9 cytoplasmic domain contains six Tyr residues (Fig 5A), and bioinformatic analyses revealed Y^980^ as a putative EGFR target site (Fig 5B). To investigate whether Y^980^ is phosphorylated, we purified full-length TLR9 from CpG-treated cells and performed quantitative mass spectrometric analyses. Mass spectrometric analyses revealed the presence of two TLR9 cytoplasmic domain-specific phospho-peptides containing pY^870^ and pY^980^ (Fig 5C). Mutation of either of these two Tyr residues (Y870F and Y980F) significantly inhibited TLR9-induced TNF levels in CpG-treated cells (Fig 5D). Mass spectrometry, followed by functional analyses, therefore, revealed phosphorylation of Y^870^ and Y^980^ was essential for TLR9-mediated gene induction.

**Fig 5.**
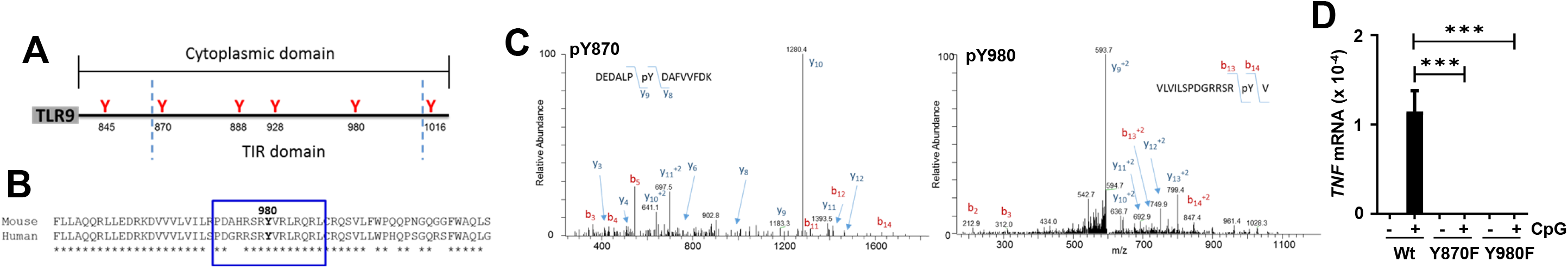
Two specific TLR9 tyrosine residues are phosphorylated after ligand stimulation. **A)** Location of tyrosines in the TLR9 cytoplasmic domain and the TIR domain. **B)** Predicted EGFR target tyrosine phosphorylation sites using GPS 6.0 in blue square. **C)** 293XL-hTLR9-HA cells were treated with CpG ODN for 30 min and then immunoprecipitated with anti-HA for TLR9 to analyze for phosphorylation of the cytoplasmic TLR9 tyrosines using LC-MS/MS. Two phospho-peptides, DEDALPpYDAFVVFDK (pY870) and VLVILSPDGRRSRpYV (pY980) were identified. The MS/MS spectra for the phosphorylated peptides are shown. The doubly charged peptide (DEDALPpYDAFVVFDK) has an observed m/z of 932.9263 Da and is within 2.63 ppm of the expected mass. The masses of the y9 and y8 ions are consistent with phosphorylation at Y870. The triply-charged peptide (VLVILSPDGRRSRpYV) has an observed m/z of 603.9943 Da and is within -2.81 ppm of the expected mass. The masses of the b14 and b13 ions are consistent with phosphorylation at Y980. **D)** 293XL-hTLR9-YFP-flag (Wt), 293XL-hTLR9-YFP-flag Y870F (Y870F), and 293XL-hTLR9-YFP-flag Y980F (Y980F) cells were treated with CpG ODN for 6 hr and induced TNF mRNA was measured by qRT-PCR.

To determine whether EGFR and Syk phosphorylate these Tyr residues, we purified TLR9 from CpG-treated Wt, EKD, or SKD cells and performed quantitative mass spectrometry. Targeted analyses revealed the relative abundance of pY^870^-containing phospho-peptide was reduced in SKD cells compared to Wt or EKD cells (Fig 6A, left panel). In contrast, pY^980^-containing phospho-peptide abundance was reduced in both EKD and SKD cells, compared to Wt cells (Fig 6A, right panel). These results indicated that Y^870^ was likely phosphorylated by Syk, and Y^980^ was likely phosphorylated by EGFR. Moreover, since EGFR was not required for Syk recruitment to TLR9 (Fig 3D), EKD cells showed pY^870^ (Fig 6A, left panel). Whereas, since Syk activity was required for EGFR activation (Fig 3E), SKD cells displayed reduced pY^980^ (Fig 6A, right panel). To determine whether phosphorylation of these Tyr residues was inter-dependent, we used the TLR9 mutants (Y870F and Y980F) and performed quantitative mass spectrometry. As expected, the Y870F mutant completely abolished the abundance of pY^870^-containing phospho-peptide without affecting pY^980^ (Fig 6B, left panel). Similarly, pY^980^ signal was completely lost in the Y980F mutant but not in Y870F (Fig 6B, right panel). Therefore, phosphorylation of the two Tyr was independent of each other. We confirmed the mass spectrometric results using biochemical approach; SKD cells, lacking both pY^870^ and pY^980^ (Fig 6A), exhibited reduced pTLR9, compared to Wt cells (Fig 6C), demonstrating Syk as a new Tyr kinase phosphorylating TLR9. We further used TLR9 mutants to evaluate whether Syk-recruitment to TLR9 required any of these two pY residues. Syk interacted with Wt as well as both Y870F and Y980F mutants of TLR9 with similar affinities (Fig 6D), indicating phosphorylation of TLR9 was not a prerequisite for Syk recruitment. TLR9-EGFR interaction, as expected, was independent of TLR9 phosphorylation; both Y870F and Y980F mutants, like Wt TLR9, interacted similarly with EGFR (Fig 6E). In the later phase, likely when TLR9 is completely phosphorylated, Syk was dissociated from TLR9 (Fig 3A). However, partially phosphorylated TLR9, in Y870F or Y980F, could retain Syk in the later phase (Fig 6E), further indicating phosphorylation of TLR9 led to Syk dissociation, presumably facilitating signalosome assembly. Together, our results demonstrated that TLR9 cytoplasmic domain was phosphorylated, sequentially, by Syk and EGFR, on Y^870^ and Y^980^, respectively, to achieve a fully functional TLR9.

**Fig 6.**
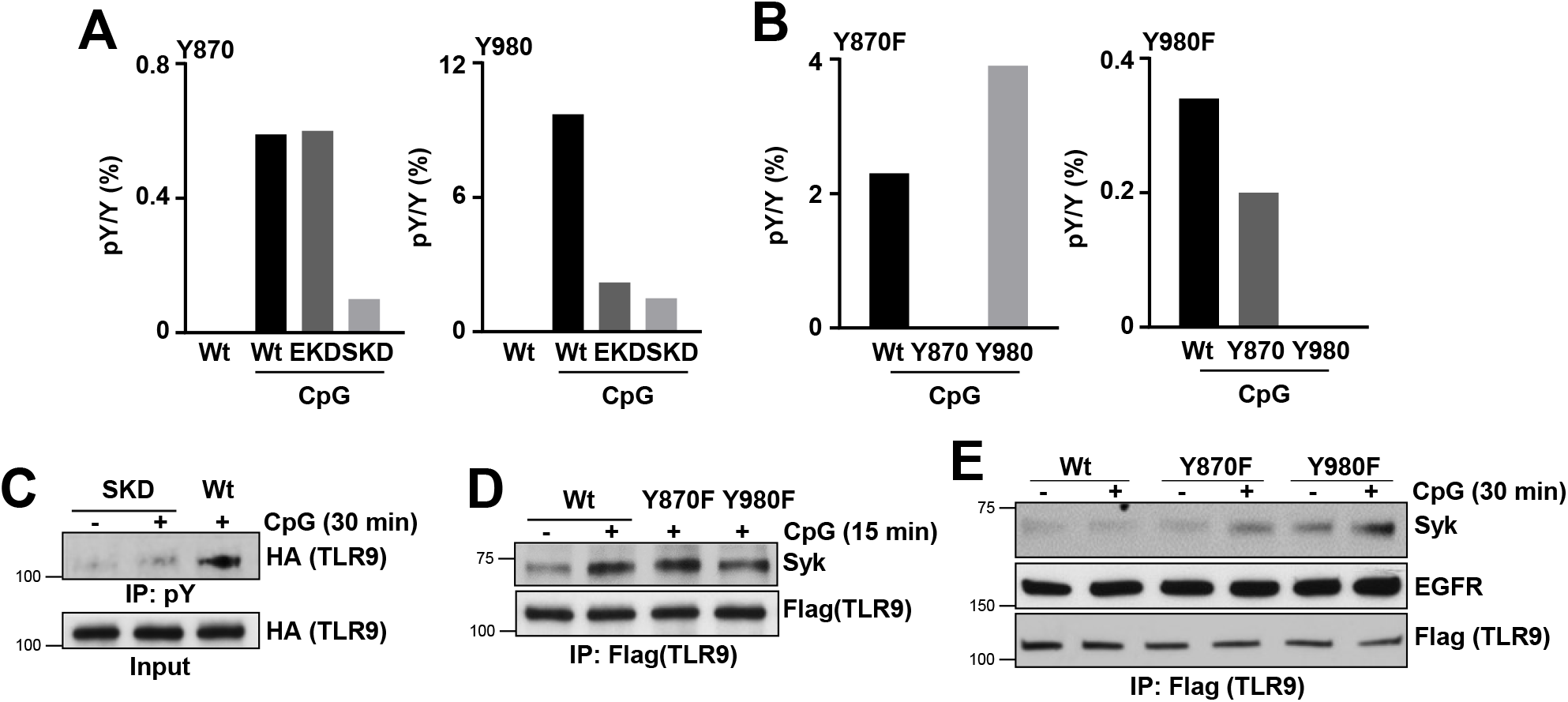
EGFR phosphorylates Y^980^ and Syk phosphorylates Y^870^ of TLR9 and the two phosphorylation events are independent of each other. **A)** Wt, shSyk (SKD), and shEGFR (EKD) 293XL-hTLR9-HA cells were treated with CpG ODN for 30 min and then immunoprecipitated with anti-HA for TLR9 to analyze the phosphorylation of cytoplasmic TLR9 tyrosines using LC-MS/MS. The left panel shows the percentage peak area ratios of phospho-peptide and unmodified peptide (pY/Y %) for Y870 and the right panel for Y980. **B)** 293XL-hTLR9-YFP-Flag (Wt), and TLR9 tyrosine mutants 293XL-hTLR9-YFP-flag Y870F (Y870F) and 293XL-hTLR9-YFP-flag Y980F (Y980F) cells were used. Treatment and analysis are as in ‘A’. The left panel shows pY/Y % for Y870 and the right panel for Y980 **C)** 293XL-hTLR9-HA cells Wt and shSyk (SKD) were treated with CpG DNA for 30 min. Cell extracts were immunoprecipitated with anti-phospho tyrosine (pY) and immunoblotted with anti-HA for TLR9. **D)** 293XL-TLR9-YFP-flag (Wt), 293XL-hTLR9-YFP-flag Y870F (Y870F), and 293XL-hTLR9-YFP-flag Y980F (Y980F) cells were treated with CpG ODN for 15 min. Cell extracts were immunoprecipitated with anti-flag for TLR9 and immunoblotted with anti-Syk and anti-flag for TLR9. **E)** The indicated cells were treated with CpG ODN for 30 min. Cell extracts were immunoprecipitated with anti-flag for TLR9 and then immunoblotted with anti-Syk, anti-EGFR, and anti-flag for TLR9.

### CpG DNA activates Syk by scavenger receptor A and Lyn, independent of TLR9

Given Syk’s essential role in phosphorylating TLR9, we investigated the mechanism of its activation in CpG-treated cells. CpG treatment caused rapid activation of Syk, analyzed by its phosphorylation on Y^352^ and Y^525^, in HEK cells (Fig 7A). Kinase activation of Syk requires Lyn-mediated pY^352^, followed by autophosphorylation on Y^525^, which was also observed in CpG-treated cells (Fig 7A). Lyn, the Syk-activating Tyr kinase, was also activated rapidly in CpG-treated HEK cells (Fig 7B). To investigate the role of TLR9 in Syk and Lyn activation, we used HEK-TLR9 cells, in which CpG treatment also caused rapid activation of Lyn and Syk (Fig 7C). Lyn activation in these cells, however, was transient; it was dephosphorylated 15 min post-CpG-treatment (Fig 7C). R406, a Syk kinase inhibitor, blocked pY^525^, indicating CpG treatment caused autophosphorylation, which is required for activation of Syk (Fig 7D). Since Syk was activated by CpG treatment in both HEK and HEK-TLR9 cells, its activation was TLR9-independent. To determine the receptor activating Lyn and Syk in the absence of TLR9, we examined scavenger receptor A (SR-A), a pattern recognition receptor on the plasma membrane, which binds CpG and activates SFKs in myeloid cells (32, 33). We used Rhein (RA), a chemical inhibitor of SR-A, to examine TLR9-independent CpG-induced pLyn and pSyk. CpG-induced pLyn and pSyk were inhibited by RA in HEK cells, indicating SR-A was involved in activation of Lyn and Syk (Fig 7E). Since SR-A was required for CpG-activated Lyn and Syk, we evaluated its role in TLR9-induced genes. RA treatment caused significant inhibition of TLR9-induced TNF and IFNB1 in HEK-TLR9 cells (Fig 7F). The role of SR-A was further evaluated in mouse myeloid cells; RA treatment also inhibited TLR9-induced Il6 and Ifnb1 in RAW264.7 macrophages (Fig 7G). Together, these results revealed a new role of SR-A in CpG-mediated activation of Syk, which is essential for phosphorylating TLR9 and downstream signaling.

**Fig 7.**
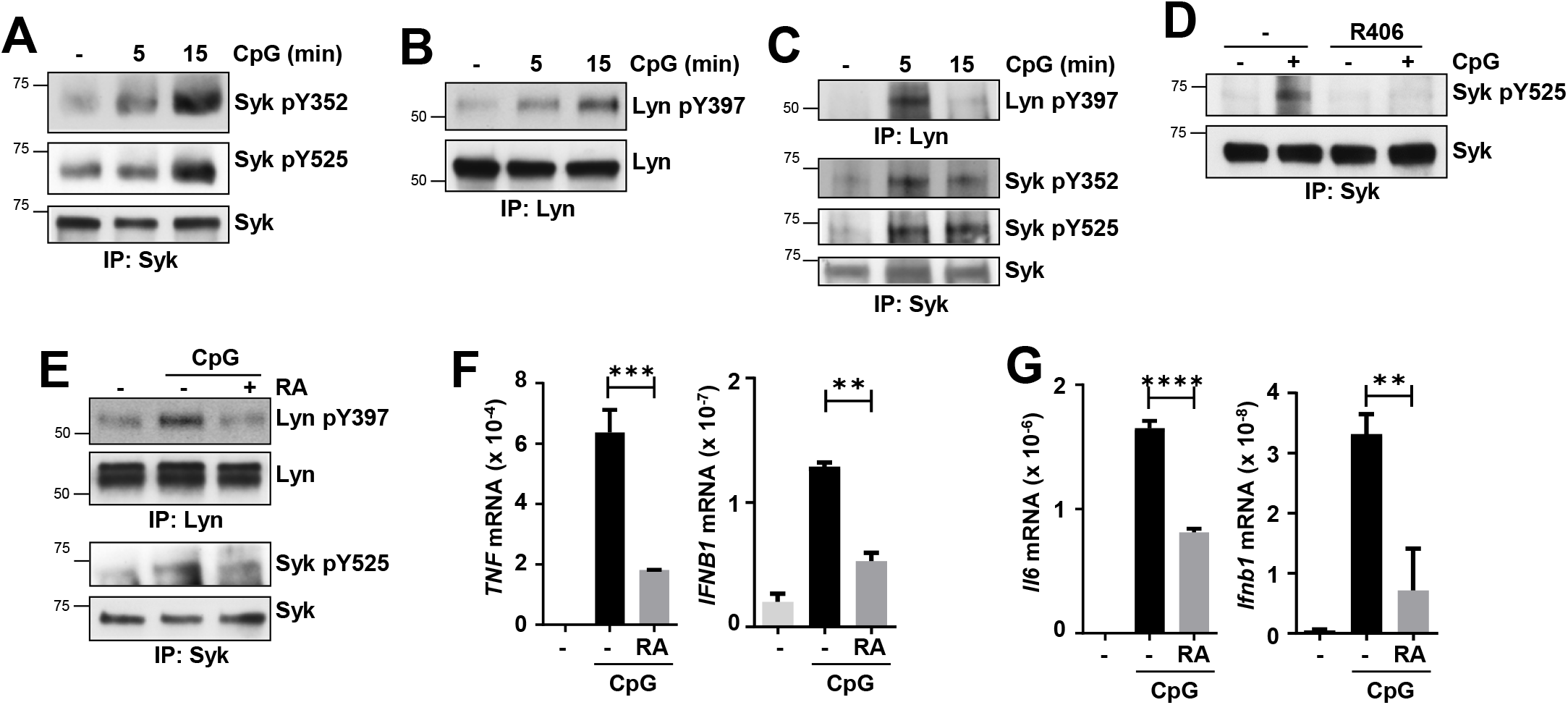
Scavenger receptor A is required for CpG-activated Syk and TLR9-induced genes. **A, B)** 293XL cells were treated with CpG DNA (10 μg/ml) for different lengths of time, and the cell extracts were subjected to immunoprecipitation (IP) followed by immunoblot as indicated. **C)** 293XL-hTLR9-HA cells were treated with CpG ODN (10 μg/ml) for the indicated time and then immunoprecipitated (IP) with anti-Lyn or anti-Syk and immunoblotted with the indicated antibodies. **D)** 293XL-hTLR9-HA cells were pre-treated for 15 min with the Syk inhibitor, R406, or the vehicle (DMSO) and then treated with CpG ODN along with R406 or DMSO for 15 min. The cell extracts were immunoprecipitated with anti-Syk and immunoblotted with anti-Syk pY525 and anti-Syk. **E)** 293XL cells were pretreated with DMSO (-) or scavenger receptor-A (SR-A) inhibitor, rhein (RA) (100 μM) for 2 hr and then treated for 15 min with CpG ODN, along with DMSO. The cell extracts were immunoprecipitated with anti-Lyn or anti-Syk and immunoblotted with anti-Lyn pY397, anti-Syk pY352, anti-Lyn and anti-Syk. **F)** 293XL-hTLR9-HA cells were pretreated with DMSO or scavenger receptor-A (SR-A) inhibitor, rhein (RA) (100 μM) for 2 hr and then treated with CpG ODN along with DMSO or RA for 6 hr and induced TNF and IFNB1 mRNAs were measured by qRT-PCR. **G)** RAW264.7 cells were treated and analyzed as in ‘E’ for induced Ifnb1 and Il6 mRNAs by qRT-PCR.

## Discussion

In this study, we reveal the molecular details of the early stages of ligand-mediated TLR9 activation (Fig 8). TLR9, like other endosomal TLRs, requires posttranslational modifications, e.g., Tyr phosphorylation of cytoplasmic domain, for its cellular functions. Previously, we showed that TLR9 binds to an endosomal pool of EGFR, a Tyr kinase, constitutively, enabling it to phosphorylate the cytoplasmic domain of TLR9 upon CpG stimulation (30). In the current study, we demonstrated that TLR9-bound EGFR required phosphorylation, an activation signal, mediated by another Tyr kinase, Syk. Syk was recruited to TLR9 upon CpG stimulation, and Syk kinase activity was essential for the activation of TLR9-bound EGFR. These results led us to postulate that TLR9 full activation required phosphorylation by EGFR and Syk. Quantitative mass spectrometric analyses revealed Syk and EGFR sequentially phosphorylated Y^870^ and Y^980^ of TLR9 cytoplasmic domain, respectively. Mutation of either of these two Tyr residues led to incomplete activation of TLR9, resulting in inhibition of TLR9-mediated gene induction. Unlike EGFR, Syk recruitment to TLR9 required its phosphorylation, mediated partially by Lyn, a cytosolic Tyr kinase. Lyn was activated, surprisingly, independent of TLR9, by a membrane-bound scavenger receptor, SR-A upon CpG binding. Together, our results led to a two-step activation model for TLR9 mediated by two Tyr kinases, Syk and EGFR.

**Fig 8.**
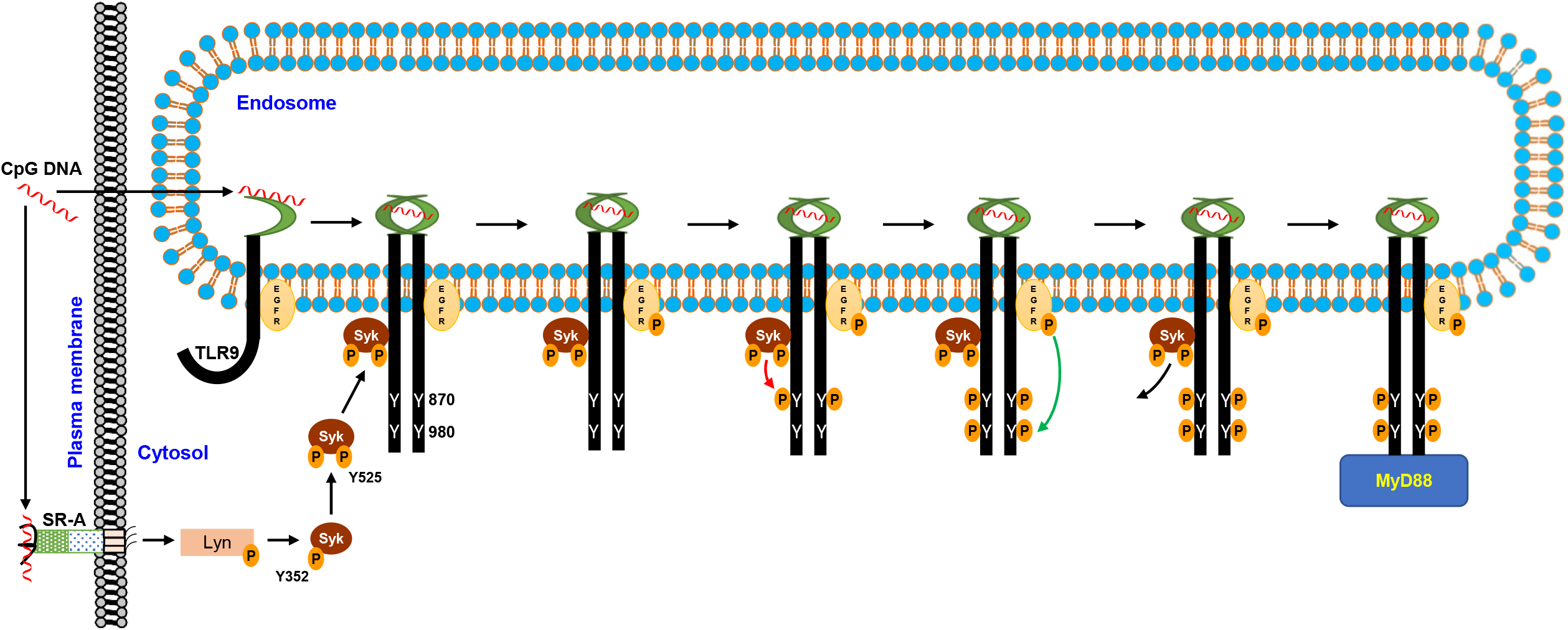
TLR9 is phosphorylated by EGFR and Syk for its activation. CpG DNA stimulation causes SR-A-mediated activation of Lyn, which phosphorylates Syk. Phosphorylated Syk (p-Syk) further autophosphorylates itself to form the activated Syk that gets recruited to endosomal TLR9, which binds EGFR constitutively. Syk recruitment to TLR9 activates EGFR by autophosphorylation; activated Syk and EGFR sequentially phosphorylate TLR9 on Y870 and Y980, respectively. Doubly phosphorylated TLR9 acts as a fully activated receptor for intracellular signaling via MyD88.

TLRs remain in closed conformation to maintain inactive states to avoid undesired activities. Activation of TLRs requires their posttranslational modifications, often mediated by Tyr phosphorylation (25). Upon binding its ligand, dsRNA, generated during viral replication or tissue damage, the cytoplasmic domain of TLR3 gets phosphorylated on two Tyr residues (29). Two Tyr kinases, EGFR and Src, phosphorylate Y^858^ and Y^759^ of TLR3 cytoplasmic domain, respectively. Phosphorylation of these residues results in the recruitment of downstream adaptor protein, TRIF, which, via a series of signaling proteins, activates IRF3 and NF-κB, to induce antiviral and inflammatory genes. In contrast, TLR4 signaling requires EGFR for activating IRF3 but not NF-κB (34). TLR9, on the other hand, required EGFR and Syk for phosphorylating Y^980^ and Y^870^, respectively, for full activation. These studies clearly indicate the EGFR requirement for endosomal TLRs. A structural role of Y^870^ for TLR9 has also been shown independent of Tyr phosphorylation (35). Mutation of Y^870^ causes immature processing of TLR9, leading to impaired downstream signaling. Recent studies further suggest that EGFR is required for cytoplasmic DNA-activated cGAS/STING signaling pathway (36, 37). Ligand-activated STING interacts with EGFR, which then phosphorylates Y^245^ of STING. STING phosphorylation is required for its intracellular trafficking, a step critical for its downstream signaling. EGFR, however, is not required for cytoplasmic RNA sensors, the RIG-I like receptors (RLRs), indicating its specificity for endosomal TLRs and STING, which are transmembrane proteins. The requirement of EGFR for TLRs and STING opens therapeutic opportunities to use FDA-approved EGFR inhibitors to dampen the undesired activation of these receptors to control viral inflammatory dysfunctions or autoimmune diseases. Polymicrobial infection, a commonly seen disease, activates multiple cellular sensors simultaneously. Under such scenarios, it needs to be examined whether EGFR can be selectively utilized by specific sensors or EGFR can help crosstalk between multiple sensors to provide innate defense.

An unexpected finding from our studies was CpG-mediated activation of Syk by SR-A and independent of TLR9. Scavenger receptors function as co-activators of multiple TLRs to help facilitate ligand binding. We report a novel crosstalk between TLR9 and SR-A mediated by Lyn and Syk. SR-A, upon recognition of CpG ODN, activated Lyn, which further led to the activation of Syk, a kinase for TLR9. The requirement of Lyn in the context of TLR9 signaling, therefore, was primarily to activate Syk, the executioner kinase. It would be interesting to see if B cells, which are major responders of TLR9 functions, might utilize BCR-activated Syk, independent of SR-A. Antigen-activated BCR recruits Syk, which may function as a TLR9 kinase for IRF and NF-κB activation. CpG-induced Syk activation has been shown before in B cells; however, this was dependent on TLR9 (38). Syk, therefore, may have cell type-specific activation mechanisms in TLR9 signaling. A role of Syk in TLR9 signaling has previously been described in fungal infection (39). Dectin-1, a receptor for fungal cell wall carbohydrates, can activate Syk, which facilitates TLR9 processing and trafficking to the phagosome. In this context, whether Dectin-1-activated Syk also phosphorylates TLR9 remains to be investigated. An anti-apoptotic function of TLR9 was observed in *L. donovani*-infected macrophages (40). *L. donovani* DNA activates TLR9, which, upon recruitment of Syk, can delay the macrophage apoptosis. Syk, therefore, can regulate various functions of TLR9 context dependently.

In addition to Syk, TLR9 itself or components of TLR9 signaling complex can be phosphorylated by other Tyr kinases cell-specifically. Bruton’s Tyr Kinase (BTK) plays a key role in TLR9 functions either by targeting TLR9 directly or proteins involved in TLR9-mediated intracellular signaling. A plasmacytoid dendritic cell (pDC) specific role of BTK has been characterized, specifically for TLR9 but not TLR7 functions (41). BTK inhibitors specifically block TLR9-mediated interferon and inflammatory responses in pDCs. BTK has been shown to mediate a synergistic function in B cells by allowing TLR9 and BCR colocalization in cellular autophagosome-like compartments (42). BTK is required for synergistic cytokine response by TLR9 and BCR in human and murine B cells, with implications in B cell activation and autoimmune diseases. DOCK8, an adaptor of the TLR9 signaling pathway in B cells, gets Tyr phosphorylated by Pyk2, and the phosphorylation is essential for its interaction with TLR9 (38). DOCK8 phosphorylation is required for TLR9-activated Src-Syk-STAT3 signaling cascade, essential for B cell proliferation and immunoglobulin production.

Although TLR9 has been studied extensively in the context of autoimmune diseases, it has implications for unexpected cellular functions. TLR9, known to recognize unmethylated CpG DNA from the microbial genome, can be activated by mitochondrial DNA (mtDNA) as well (14, 43, 44). In dengue virus (DenV)-infected human dendritic cells, mtDNA can activate TLR9 to induce antiviral gene expression (45). The antiviral role of TLR9 against RNA viruses is surprising and opens up possibilities for investigating TLR9 functions in RNA virus infections. The mtDNA as a cellular sensor for TLR9 has been investigated in a number of cellular functions and disease conditions. In cancer stem cells, mitophagy-derived mtDNA activates TLR9, which promotes Notch1 activity. TLR9-activated Notch1 further promotes AMPK activity, which supports tumor cell growth (46). A pathophysiologic role of TLR9 has been shown in non-alcoholic steatohepatitis (NASH) via recognition of mtDNA (22). TLR9, expressed in myeloid cells, gets activated by hepatocyte-derived mtDNA in promoting NASH. TLR9 activation by mtDNA can lead to long-term depression, and this function is associated with TLR9-mediated regulation of neuronal activity (14). In platelets, carboxyalkylpyrrole protein adducts (CAPs), generated during oxidative stress, can serve as physiological ligands of TLR9 in platelets (47). Endogenous ligand, CAPs activate TLR9/MyD88/IRAK1 signaling pathway and result in platelet activation and thrombosis. These studies provide opportunities to use TLR9 antagonists for therapeutic interventions. Our study, which adds EGFR, Syk, and Lyn in TLR9 functions, may provide unique opportunities to examine the clinical inhibitors of these kinases in TLR9-induced disease pathologies.

### Experimental Procedures

#### Reagents and antibodies

Syk inhibitor (inh-r406), R406 was from Invivogen, Gefitinib (S1025) and Scavenger receptor A inhibitor (Rhein, s2400) were obtained from Selleckchem, and CpG oligonucleotide (CpG-B) was from Integrated DNA Technologies (IDT). Phospho-tyrosine antibody (05-321) was obtained from Millipore, antibodies against EGFR (2646), phospho-EGFR Y1068 (2234), MyD88 (4283), Syk (13198), phospho-Syk Y352 (2701), phospho-Syk Y525 (2711), Lyn (2796), and actin (A5441) were from Cell Signaling Technology, anti-phospho Lyn Y397 (orb315590) was from Biorbyt, HA antibody (ab18181) was from Abcam, Chloroquine (C6628), anti-flag M2 affinity gel (A2220), and anti-flag antibody (F7425) were from Sigma Aldrich, anti-YFP (SC-32897) and anti-Scavenger receptor-A (SR-A, SC-166184) from Santa Cruz Biotechnology.

#### Cell lines

RAW264.7 cells from ATCC were maintained in DMEM containing 10% FBS and penicillin-streptomycin. 293XL and 293XL-human TLR9-HA cells from Invivogen were maintained in DMEM containing 10% FBS, penicillin-streptomycin, normocin, and blasticidin as per company instruction. 293XL and HT1080 cells with human TLR9 (with YFP and Flag double epitope tags), TLR9 Y870F and Y980F were generated using lentiviral transduction. EGFR-knockdown, Syk-knockdown, and Lyn-knockdown in 293XL-human TLR9-HA cells were generated by lentiviral transduction of human EGFR-specific shRNA (Sigma, #TRCN0000121202), Syk-specific ShRNA (Sigma, #TRCN0000197242), and Lyn-specific ShRNA (Sigma, # TRCN0000218210) respectively, followed by selection under puromycin and expressions were confirmed using western blot (WB). A non-targeting shRNA (#SHC002) was used as a control. The CRISPR-induced genomic deletion was performed by overnight lentivirus transduction of subconfluent HT1080 cells LentiCRISPRv2 expressing Cas9 endonuclease and Human EGFR sgRNA sequences TGAGCTTGTTACTCGTGCCT and GAGTAACAAGCTCACGCAGT were used (48). The knockout cells were then selected under puromycin. Single cell clones were screened using genomic DNA sequencing and WB. Extracellular domain deleted EGFR (ΔExt) expression vector was kindly provided by Xiaoxia Li (49).

#### Quantitative real-time PCR

Total cellular RNA was extracted from cells using Roche RNA isolation kit (11828665001, Roche), and used for cDNA preparation using ImprompII Reverse Transcription Kit (A3803, Promega). The cDNA (0.5 ng) was applied to a 384-well plate for real-time PCR using SYBR Green PCR mix (Applied Biosystem’s Power) in Roche Light Cycler 480 II. The expression levels of the induced mRNAs were normalized to 18S rRNA or RPL32 mRNA. GraphPad Prism was used for plotting.

#### Immunoprecipitation and western blot

Cells were lysed on ice using 50 mM Tris–HCl pH 7.4 containing 150 mM NaCl, 5 mM EDTA, 1 mM DTT, 1 mM PMSF, and cocktail (Roche) for western blot and 20 mM HEPES pH 7.5 containing 150 mM NaCl, 10 mM NaF, 1.5 mM MgCl2, 10 mM β-glycerophosphate, 2 mM EGTA, 1 mM sodium orthovanadate, 0.5% (v/v) Triton X-100, and protease inhibitors (Roche Applied Science, Indianapolis, IN, USA). The lysates were pre-cleared with A/G agarose beads. The pre-cleared lysates were then treated with antibodies of targeted proteins for two hours at 4 °C. These were then incubated with A/G agarose beads (426535, Santa Cruz Biotechnology) overnight at 4 °C. The beads obtained after centrifugation were washed thrice with lysis buffer and boiled with SDS-page buffer to elute the proteins. The eluted samples were then analyzed by SDS-PAGE followed by Western blot. The samples were run through the SDS-PAGE gels and then transferred to the polyvinylidene difluoride (PVDF) membrane (Bio-Rad). The membranes were then incubated for 1 hr in 5% skim milk in TBST buffer (150 mM NaCl; Tris, pH 7.4; and 0.1% Tween 20) at room temperature. The membranes were then incubated with the primary antibody in a cold room overnight followed by the secondary HRP antibody. The membranes were then visualized by Super-Signal West Pico chemiluminescent substrate (Pierce Chemical).

#### Mass spectrometry

Finnigan LTQ-Orbitrap Elite hybrid mass spectrometer was used for LC-MS study for hTLR9-HA cells with Dionex 15 cm × 75 μm id Acclaim PepMap C18, 2 μm, 100 Å reversed-phase capillary chromatography column. The micro-electrospray ion source was operated at 2.5 kV. After treatment or non-treatment, the cells were lysed, pulled down with anti-HA or anti-flag, and then separated using SDS-PAGE. Protein digestions were done using the TLR9 bands from the gels, after washing and de-stained in 50% ethanol containing 5% acetic acid the gel pieces were dehydrated in acetonitrile, dried in a Speed-vac, and digested by adding 5 μl 10 ng/μl of trypsin, alpha-lytic or trypsin-AspN, in 50 mM ammonium bicarbonate. After overnight incubation, the peptides were extracted into two portions of 30 μl each 50% acetonitrile and 5% formic acid. The combined extracts were then evaporated to ∼ 10 μl in a Speed-vac and re-suspended in 1% acetic acid to make up to 30 μl for analysis. 5 μl of the extract was injected into the LC-MS system and the peptides eluted from the column, using an acetonitrile/0.1% formic acid gradient at a flow rate of 0.25 μl/min, were introduced into the source of the mass spectrometer online. The digest was analyzed in both a survey manner and a targeted manner. The survey experiments were performed using the data-dependent multitask capability of the instrument, acquiring full scan mass spectra to determine peptide molecular weights and product ion spectra to determine amino acid sequences in successive instrument scans. The LC-MS/MS data were then searched using Mascot and Sequest programs against the full human reference sequence database, specifically against the sequence of TLR9. The parameters used in this search include a peptide mass accuracy of 10 ppm, fragment ion mass accuracy of 0.6 Da, carbamidomethylated cysteines as a constant modification, and oxidized methionine and phosphorylation at S, T, and Y as a dynamic modification. The results were then filtered based on Mascot ion scores and Sequest XCorr scores. All positively identified phosphopeptides were manually validated. The targeted experiments involve the analysis of specific TLR9 peptides, including the phosphorylated and unmodified forms of the Y845, Y870, Y888, Y928, Y980, and Y1021 peptides. The chromatograms for these peptides were then plotted based on known fragmentation patterns. The peak areas of these chromatograms were used to determine the extent of phosphorylation (50, 51).

hTLR9-YFP samples were analyzed by LC-MS using a Fusion Lumos Tribrid MS (Thermo Scientific) equipped with a Dionex Ultimate 3000 nano UHPLC system, and a Dionex (25 cm x 75 μm id) Acclaim Pepmap C18, 2-μm, 100-Å reversed-phase capillary chromatography column. Peptide digests (5 μl) were injected onto the reverse phase column and eluted at a flow rate of 0.3 μl/min using mobile phase A (0.1% formic acid in H2O) and B (0.1% formic acid in acetonitrile). The gradient was held at 2%B for 5 minutes, %B was increased linearly to 35% in 80 minutes, increased linearly to 90% B in 10 minutes, and maintained at 90% B for 5 minutes. The mass spectrometer was operated in a data-dependent manner which involved full scan MS1 (375-1700 Da) acquisition in the Orbitrap MS at a resolution of 120000. This was followed by CID (1.6 Da isolation window) at 35% CE and ion trap detection. MS/MS spectra were acquired for 3 seconds. The second method was used for glycopeptide identification and involved full scan MS1 7 (350-1700 Da) acquisition in the Orbitrap MS at a resolution of 120000. Dynamic exclusion was enabled where ions within 10 ppm were excluded for 60 seconds.

#### Confocal microscopy

EGFR KO HT1080-TLR9 (with YFP and Flag double epitope tags) cells, reconstituted with extracellular domain deleted EGFR, were grown on glass coverslips, and the cells were fixed at 20 min each with 4% paraformaldehyde and permeabilized with 0.2% Triton X-100. Fixed cells were blocked with 5% normal goat serum for 1 hr and then labeled overnight with anti-EEA1 (610457, BD Transduction Labs) to stain early endosomes and anti-EGFR (2646, Cell Signaling). Goat anti-mouse Alexa Fluor 647 (A32728, Invitrogen) and anti-rabbit Alexa Fluor 594 (A32740, Invitrogen) (for 1 hr) were used, respectively, as secondary antibodies. These were then mounted using VECTASHIELD-DAPI. The images were taken by confocal laser scanning microscopy (Leica TCS SP8) and were processed with Leica LCS software. ImageJ software was used for determining the co-localization of two proteins.

#### Statistical analysis

GraphPad Prism 9 software was used for all statistical analyses. Technical replicates were used to represent the qRT-PCR results and shown representatives of three independent experiments. P values were calculated using one-way ANOVA.

## Acknowledgments

This article is dedicated to celebrating the illustrious scientific career of Ganes Sen on his retirement for his seminal contributions to discoveries in innate immune signaling and cellular interferon responses. In addition to leading a robust research program, he inspired generations of researchers to follow his path of finding answers to complex biological questions. This work was supported in part by the National Institutes of Health grants CA068782 (G.C.S.), CA062220 (G.C.S), 1S10OD023436-01 (B.W.), AI155545 (S.C.), and AI165521 (S.C.). We thank Xiaoxia Li for providing the extracellular domain-deleted EGFR expression vector.

## Data Availability

All data presented in this paper are contained within the manuscript.

## Author contributions

MV, CW, BW, and GCS designed the experiments, MV, CW, PMK, and BW performed experiments and analyzed the results, MV, GCS and SC wrote and edited the manuscript.

